# Larvae and adults exhibit contrasting patterns of immune gene expression and infection resistance in wild flour beetle populations

**DOI:** 10.1101/2021.02.24.432795

**Authors:** Ann T. Tate, Abby Perry, Derrick G. Jent

## Abstract

In nature, hosts face shifting patterns of parasite exposure and life history trade-offs as they develop from birth to old age. As a result, the net fitness benefit of immunological investment can change dramatically from one life stage to the next. Previous work has revealed a puzzling diversity of relative immune investment patterns among juvenile and adult stages, and it is not clear whether lessons learned from one particular population or species can be generalized to wild populations, after accounting for local adaptation and other variance-generating processes. In this study, we quantify larval and adult immune gene expression and resistance to bacterial infection in two flour beetle species (*Tribolium castaneum* and *T. confusum*) from two lab-adapted and five wild-derived populations. Our results provide a clear signal of higher infection-induced immunological investment and resistance in adults relative to larvae, despite variation among species in immune gene regulation. Better characterization of stage-specific investment in infection resistance in natural populations can inform our understanding of life history evolution and improve predictions of disease dynamics in the wild.

## Introduction

The net contribution of different life history traits to host fitness changes across the ontogeny of the organism. While juveniles derive benefit from growing quickly to avoid predators and investing in survival to reach reproductive age, adults maximize fitness by finding mates and producing as many high-quality offspring as possible [1]. In many organisms, juveniles and adults also occupy different niches, such that traits beneficial in one life stage may not maximize fitness in another [2]. For organisms that undergo metamorphosis, such as holometabolous insects and amphibians, the dramatic reinvention of physiology can provide an opportunity to adaptively decouple juvenile and adult traits to better optimize fitness [3, 4] in stage-specific environments.

Even when life stages share a niche, juveniles and adults may face different hazards and trade-offs among life history traits [5, 6] that might favor trait divergence over evolutionary time. For example, juveniles may feed at different rates or from a different distribution of resources than adults, altering patterns of exposure to parasites, pathogens, and symbionts over host age [7–9]. Investing in immunological resistance to those parasites, however, can tax resource allocation to traits like growth and reproduction [6], contributing to asymmetry in the net cost of immunity to host fitness across different life stages [10].

Collectively, previous research on infection susceptibility and immunity in holometabolous insects defies neat conclusions about relative immune investment across ontogeny. In honeybee workers (*Apis mellifera*), for example, the activity of the melanization enzyme phenoloxidase increases steadily over development from larva to adult [11], while in the tobacco hornworm (*Manduca sexta*), melanization activity peaks in the larval stage and declines thereafter [12] (see [13] for further examples). Most studies, however, employ insect populations reared in (and evolving under) lab conditions, sometimes for substantial periods of time. After accounting for the effects of local adaptation, can we still discern generalizable principles of stage-structured immunological investment?

To address this question, we turned to wild and lab-derived populations of flour beetles (*Tribolium castaneum* and *T. confusum*). Previous research in lab-reared *T. castaneum* has suggested that the regulation of immune [13] and cuticular [14] gene expression differs among life stages in uninfected individuals, while, in larvae specifically, different populations of both species exhibit natural variation in immune gene expression and resistance to bacterial infection [15]. To test whether immune gene expression and infection resistance in these populations differs among life stages, we injected larvae and adults from seven populations (four *T. castaneum*, three *T. confusum*) with one of four increasing doses of the entomopathogenic bacterium *Bacillus thuringiensis* (Bt) or sterile media as a wounding control. We quantified relative bacterial load and host expression of several immune genes, including the antimicrobial peptides *defensin-1 and attacin-1*, the recognition protein*pgrp-sc2*, and the melanization enzyme *ddc*, during the acute phase of infection. Our results highlight the importance of life stage for predicting disease susceptibility and underscore the complex relationships between development, phylogeny, and ecology in life history evolution.

## Methods

### Experimental Design

The initial collection of wild beetle populations and analysis of variation in immune gene expression among larvae from these populations has been described in detail in [15]. The current study compares these larval data (available from Data Dryad [16]) to data from adults processed at the same time (data available i). Briefly, we created breeding groups of 18 adult beetles (~9 male and 9 female) from two lab stock colonies (Snavely *T. confusum* and Snavely *T. castaneum*) and five wild-derived, parasite-free stock colonies (Green River *T. castaneum*, Green River *T. confusum*, Marshall *T. confusum*, Dorris *T. castaneum*, WF Ware *T. castaneum*). Breeding groups were allowed to lay eggs in whole wheat flour before being passaged to new flour every 48 hours. The adult cohorts from this experiment were derived from the second egglay, while the larvae were derived from the ninth egglay. We have previously determined that egg laying order within the first few weeks minimally affects Bt resistance traits in flour beetles [17]. Individuals from the different breeding groups were randomly assigned to different initial doses immediately before infection. We kept track of sex for a subset of the adults (N = 45) derived from the Dorris, Green River, and WF Ware *T. castaneum* populations to determine whether adult sex could account for variation in gene expression. As preliminary analyses did not find a strong effect, however, and not all adults (or any of the larvae) were classified by sex, we dropped this variable from subsequent analyses.

An overnight culture of *Bacillus thuringiensis* Berliner (ATCC 55177) was prepared from a glycerol stock and grown at 30°C in Nutrient Broth #3 (NB, Sigma Aldrich). An aliquot of overnight culture was brought to an OD of 0.5 in fresh NB, and initial doses were created by mixing 500uL overnight culture with 500uL log phase culture (~1*10^8^ CFU/mL) and then serially diluting aliquots with sterile NB to create the desired two-fold dilution. The second-highest dose is the larval LD50.

When the larval cohorts reached approximately 4mm in length (about 3 days before the onset of pupation), we injected both larvae and unmated adults (approximately 7 days post eclosion) with either sterile NB or one of four two-fold increasing doses of Bt (N = 6-8 individuals per dose per stage per population), using a micropin dipped in the relevant solution. All injected beetles were sacrificed at 8 hours post injection by flash freezing, and stored at −80°C. We extracted RNA, created cDNA, and quantified gene expression via RT-qPCR as previously described [15] using degenerate primers capable of amplifying transcripts from both flour beetle species with equal efficiency for four immune genes: *attacin-1, defensin-1,pgrp-sc2*, and *ddc*. In addition, we quantified the expression of host reference gene *rps18* and a fragment of the Bt 16s gene that has been previously validated as a proxy for bacterial CFU [17]. We calculated ΔCt values (target gene Ct – reference gene Ct) for each gene template for each individual. These values were linearized (2^-ΔCt^) prior to analysis and then log-transformed for normality, as in [18, 19].

### Statistical Analysis

To analyze the effect of life stage on resistance at 8 hours post infection, we employed linear mixed models of the form: relative bacterial load ~ life stage + initial dose + (1ļpopulation), where stage and dose are main effects and population is treated as a random effect in the model. To investigate whether life stage affects the magnitude of inducible immunity to a wounding event or infection, we analyzed the relative magnitude of gene expression at eight hours post exposure. For each analyzed gene, the linear mixed model took the form of: gene expression ~ stage + (1 ļpopulation) for the sterile stabs and gene expression ~ stage + log2(bacterial load) + (1 ļpopulation) for the infected individuals. Species was added as an additional fixed effect if it significantly improved model fit, as determined by a log likelihood ratio test.

Because p-values cannot be reliably calculated for mixed models, we instead rely on confidence intervals (2.5%, 97.5%) to determine significance, where a lack of overlap with 0 encourages the rejection of the null hypothesis. All statistical analyses were conducted using the lmer function from the lme4 package in R (v.4.0.3).

## Results

### Impact of life stage on resistance and immune gene expression

Both initial dose and life stage significantly predicted relative bacterial load at 8 hours post infection (**Table 1**). The bacterial load increased with initial dose, and larval bacterial load was about 1.5 times greater than adult bacterial load at 8 hours post infection (**Fig. 1**).

**Table 1.** The effect of life stage on bacterial load at 8 hours post injection.

|  | Estimate | Std. Error | t value | 2.5CI | 97.5CI |
| --- | --- | --- | --- | --- | --- |
| Intercept | -14.73 | 0.66 | -22.18 | -16.06 | -13.40 |
| Initial dose | 0.86 | 0.09 | 9.33 | **0.68** | **1.04** |
| Stage | -0.62 | 0.20 | -3.05 | **-1.02** | **-0.22** |

Notes: Model is $\log_2(\text{bacterial load}) \sim \text{dose} + \text{stage} + (1|\text{pop})$ . Larvae are the baseline stage. Adding species as a factor did not improve model fit (as determined by log likelihood ratio test) and was excluded from the model. Bold confidence intervals indicate lack of overlap with null expectation.

**Figure 1.**
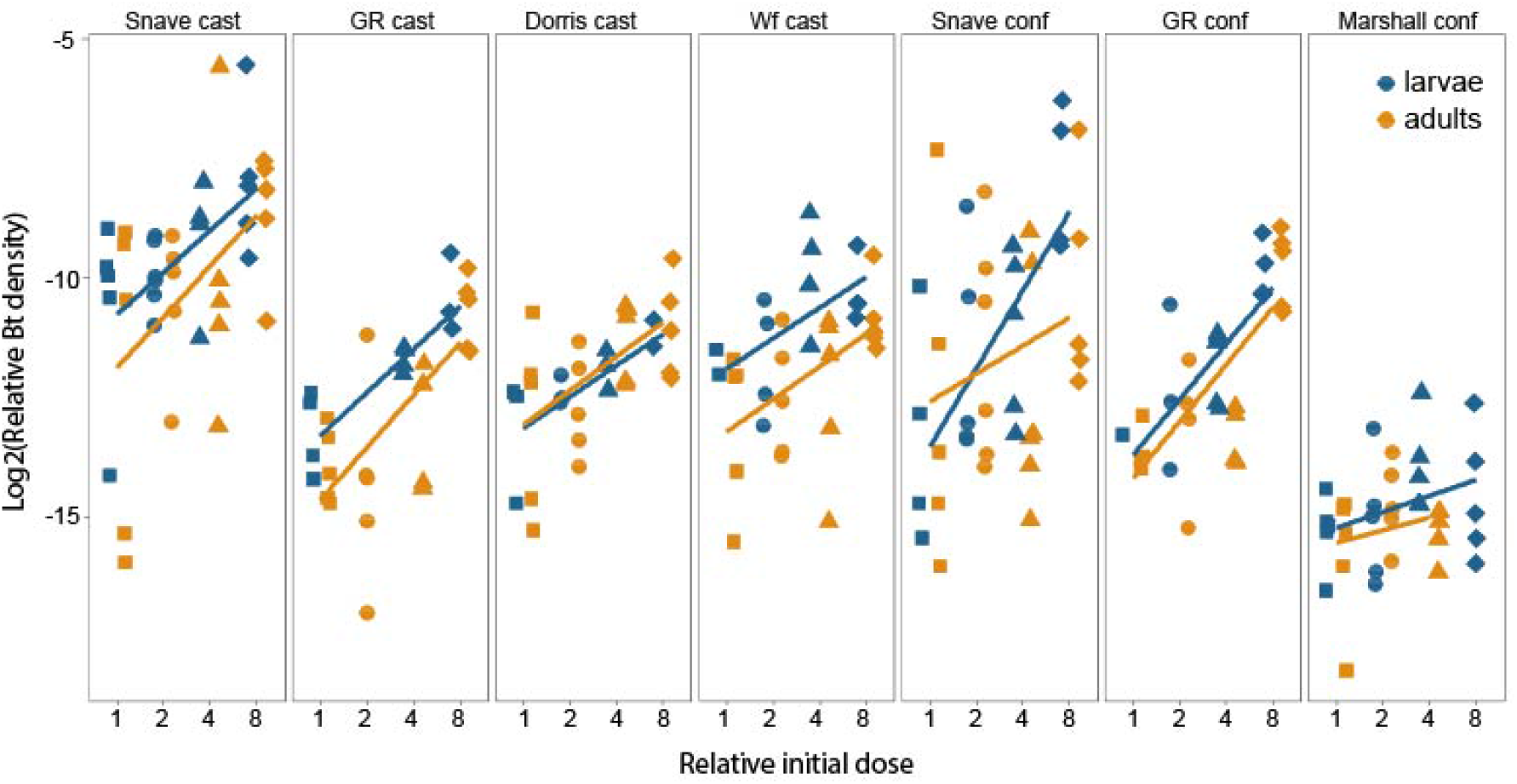
The impact of life stage on bacterial load in seven flour beetle populations. Larvae (blue) and adults (orange) were infected with one of four two-fold increasing doses (1,2,4,8) and sacrificed at 8 hours post infection. Bacterial load was estimated via RT-qPCR relative to host reference gene expression. Shapes reflect initial dose. Cast = *T. castaneum*, conf = *T. confusum*.

After injection with sterile media, life stage did not significantly predict expression of any gene (**Table 1**). The expression of *defensin-1* by different life stages across populations is depicted in **Fig. 2A**. While model selection did not generally favor the inclusion of species for most genes, the effect of species on *attacin-1* was significant, with *T. confusum* exhibiting higher expression than *T. castaneum*.

**Figure 2.**
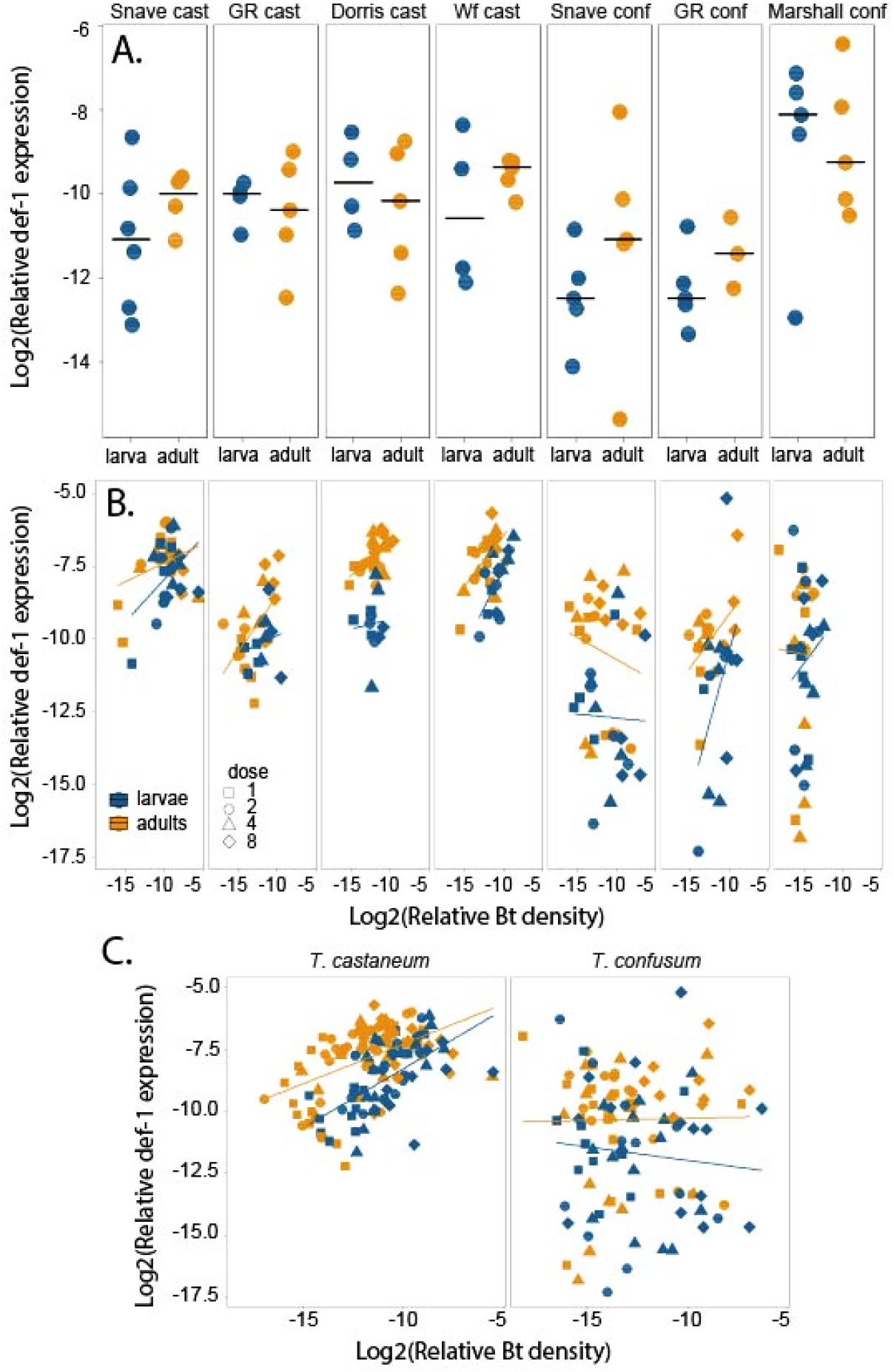
The impact of life stage on inducible expression of the antimicrobial peptide *defensin-1* in seven flour beetle populations. Larvae (blue) and adults (orange) were treated with either a sterile media jab (**A.**) or infected with one of four two-fold increasing doses (**B**) and sacrificed at 8 hours post infection. *Defensin-1* expression and bacterial load were estimated via RT-qPCR relative to host reference gene expression; aggregated infection-induced expression data by species is visualized in (**C**). Shapes in (**B, C**) reflect initial dose while log2(relative Bt density) represents relative bacterial density at 8 hours. Cast = *T. castaneum*, conf = *T. confusum*.

At 8 hours post injection with live bacteria, adults exhibited significantly higher expression of all immune genes (**Table 2**), which was relatively consistent across populations (**Fig. 2B**). The antimicrobial peptide *defensin-1* exhibited the biggest effect size between stages after accounting for variance among populations (**Fig. 2C**). As expected, the main effect of relative bacterial load on expression was significant for all genes. The main effect of species was significant for *defensin-1* and *ddc* only, where *T. confusum* exhibited lower expression of both genes relative to *T. castaneum* (**Fig. 2C**).

**Table 2.** The effect of life stage on immune gene expression at 8 hours post injection with sterile saline or Bt.

|  | Estimate | Std. error | t value | 2.5CI | 97.5CI |
| --- | --- | --- | --- | --- | --- |
| *Attacin-1 (sterile)* |  |  |  |  |  |
| Intercept | -13.33 | 0.48 | -27.98 | -14.37 | -12.28 |
| Stage | -0.07 | 0.38 | -0.20 | -0.82 | 0.68 |
| Species | 1.57 | 0.67 | 2.35 | **0.03** | **3.07** |
| *Defensin-1 (sterile)* |  |  |  |  |  |
| Intercept | -10.75 | 0.38 | -28.01 | -11.58 | -9.91 |
| Stage | 0.61 | 0.35 | 1.75 | -0.08 | 1.32 |
| *Pgrp-sc2 (sterile)* |  |  |  |  |  |
| Intercept | -11.62 | 0.65 | -17.76 | -13.09 | -10.15 |
| Stage | 0.37 | 0.23 | 1.63 | -0.08 | 0.82 |
| *Ddc (sterile)* |  |  |  |  |  |
| Intercept | -11.44 | 0.20 | -57.38 | -11.84 | -11.01 |
| Stage | 0.56 | 0.27 | 2.06 | -0.02 | 1.11 |
| *Attacin-1 (infected)* |  |  |  |  |  |
| Intercept | -6.51 | 0.73 | -8.92 | -7.87 | -5.15 |
| Bt load | 0.34 | 0.04 | 7.68 | **0.26** | **0.43** |
| Stage | 0.81 | 0.17 | 4.86 | **0.48** | **1.13** |
| Species | -0.60 | 0.81 | -0.74 | -2.14 | 0.94 |
| *Defensin-1 (infected)* |  |  |  |  |  |
| Intercept | -7.00 | 0.85 | -8.27 | -8.59 | -5.41 |
| Bt load | 0.17 | 0.06 | 2.77 | **0.05** | **0.29** |
| Stage | 1.28 | 0.24 | 5.33 | **0.80** | **1.74** |
| Species | -2.40 | 0.70 | -3.42 | **-3.73** | **-1.06** |
| *Pgrp-sc2 (infected)* |  |  |  |  |  |
| Intercept | -8.73 | 0.92 | -9.45 | -10.69 | -6.72 |
| Bt load | 0.14 | 0.04 | 3.51 | **0.06** | **0.22** |
| Stage | 0.38 | 0.15 | 2.57 | **0.09** | **0.67** |
| Species | -0.40 | 1.24 | -0.32 | -3.19 | 2.40 |
| *Ddc (infected)* |  |  |  |  |  |
| Intercept | -6.60 | 0.47 | -14.15 | -7.53 | -5.61 |
| Bt load | 0.30 | 0.04 | 8.04 | **0.22** | **0.37** |
| Stage | 0.55 | 0.14 | 3.80 | **0.26** | **0.83** |
| Species | -1.28 | 0.31 | -4.07 | **-1.97** | **-0.55** |

Notes: Linear mixed models (using maximum likelihood method) are of the general structure: gene expression ~ log2(Bt load) + stage + (1|pop). Model selection was conducted with a log likelihood ratio test to determine inclusion of species as an additional fixed effect. For infected analyses, all genes either had no difference among models with and without species, or were improved with the inclusion of species, so species was retained for all. Bold confidence intervals indicate lack of overlap null expectation.

## Discussion

Theory predicts that evolutionarily optimal levels of juvenile investment in immunological resistance depend on the precise magnitude of life history parameters like life span and juvenile background mortality rate [10], while changes in parasite exposure across stages likely also contribute to ontogenetic variation in immunological investment [20]. Since natural populations may experience very different selective pressures from parasites, predators, and other ecological factors that influence these parameters, it is difficult to predict *a priori* whether decoupling of immunological traits across life stages will show consistent patterns in the wild. Our results provide a clear signal of life stage-specific variation in resistance-associated traits to a particular infection: adult flour beetles invest more in the inducible expression of all assayed immune genes and reap a modest but detectable benefit with regard to resisting bacterial proliferation.

Is the observed stage-specific variation actually adaptive, in the sense that it maximizes the fitness of an individual by independently optimizing the response in each life stage, or could it simply reflect a developmental constraint on the deployment of the larval immune response? The dominance of the adult response to infection does not appear to extend to the wounding response, as any differences in gene expression among life stages appear to be population-specific and exhibit no consistent trend (e.g. Fig. 2A). Thus, there does not appear to be any fundamental developmental constraint manifest in the baseline inducible response. In the absence of studies that manipulate the expression of key genes independently in each life stage and measure associated fitness effects, however, it is difficult to obtain a direct answer about the relative roles of adaptation and constraint in ontogenetic decoupling of immunological phenotypes.

When evaluating the potential role of local adaptation in structuring immunological variation across stages, it is interesting to consider differences in variance structure among populations and species [21, 22]. In our study, all four populations of *T. castaneum* tell a similar story with regard to infection-induced expression of *defensin-1:* expression depends on bacterial load, and adults exhibit higher expression levels (Fig. 2C). The expression of *defensin-1* in *T. confusum* appears to be under different regulatory control since it does not depend neatly on bacterial density, but even here adults express these genes at higher levels. These two species diverged as long as 60 million years ago [23] and occupied different continents, but re-converged at the dawn of domestication to specialize on agricultural products; they even co-occur in some locations (including two in this study). Whether ontogenetic decoupling of immunity is ancestral to these beetles, or whether it arose more recently as ecological conditions converged, remains an interesting question for future study in this and other host taxa.

In addition to the implications of ontogenetic decoupling for immune system evolution, variation in resistance among life stages can influence disease dynamics in stage-structured host populations. If less resistant juvenile stages are more competent at transmitting microbes, for example, populations that skew toward more susceptible stages could drive up the force of infection [24, 25]. If, on the other hand, adult investment in immunity trades off with reproduction [26], then a decrease in the birth of new susceptibles could drive down disease prevalence even further than the effect of adult resistance alone. The relative consistency of life stage differences in immunological resistance among populations in our study lends support to the general applicability of stage-structured models parameterized to reflect immunological asymmetry.

In summary, our study provides evidence of stage-structured variation in immune responses across natural populations and species. These observations raise the question of whether these differences evolved adaptively against stage-specific parasite pressures, or whether life history trade-offs or developmental constraints dominate the optimization of immune system investment. Future work investigating the mechanistic basis of immune system regulation across host development and in response to the natural parasite milieu encountered by each life stage could help to disentangle the ecological and evolutionary contributors to ontogenetic decoupling of immune responses.

## Acknowledgements

This work was supported in part by NSF DEB grant no. 1753982 to A.T.T.

## Author Contributions

A.T.T. conceived and designed the study and experiments. A.P. and D.J. conducted the experiments. A.T.T. and A.P. analyzed the data. A.T.T. wrote the manuscript.

## Data Accessibility

Larval gene expression data can be accessed in Data Dryad [16]. Adult gene expression data can be accessed in the supplemental material.

